# Novel fragile X syndrome 2D and 3D brain models based on human isogenic FMRP-KO iPSCs

**DOI:** 10.1101/2020.11.12.379800

**Authors:** Carlo Brighi, Federico Salaris, Alessandro Soloperto, Federica Cordella, Silvia Ghirga, Valeria de Turris, Maria Rosito, Pier Francesca Porceddu, Angelo Reggiani, Alessandro Rosa, Silvia Di Angelantonio

## Abstract

Fragile X syndrome (FXS) is a neurodevelopmental disorder, characterized by intellectual disability and sensory deficits, caused by epigenetic silencing of the *FMR1* gene and subsequent loss of its protein product, fragile X mental retardation protein (FMRP). Delays in synaptic and neuronal development in the cortex have been reported in FXS mouse models, however, the main goal of translating lab research into pharmacological treatments in clinical trials has been so far largely unsuccessful, leaving FXS a still incurable disease. Here, we generated 2D and 3D in vitro human FXS model systems based on isogenic *FMR1* knock-out mutant and wild-type human induced pluripotent stem cell (hiPSC) lines. Phenotypical and functional characterization of cortical neurons derived from FMRP-deficient hiPSCs display altered gene expression and impaired differentiation when compared with the healthy counterpart. FXS cortical cultures show increased proliferation of GFAP positive cells, likely astrocytes, increased spontaneous network activity and depolarizing GABAergic transmission. Cortical brain organoid models show increased proliferation of glial cells, and bigger organoid size. Our findings demonstrate that FMRP is required to correctly support neuronal and glial cell proliferation, and to set the correct excitation/inhibition ratio in human brain development.

## INTRODUCTION

Fragile X syndrome (FXS), first described by J. Purdon Martin and Julia Bell in 1943 as a mental defect following an X-linked inheritance pattern [1], represents the second cause of inherited intellectual disability after the Down syndrome, and the most prevalent one in males [2,3]. FXS patients are characterized by cognitive impairment, defective communication, abnormal sensory reactivity, anxiety, hyperactivity, gaze aversion, and impulsivity [4]. The etiology of FXS lies in an abnormal trinucleotide repeat (CGG) expansion in the 5’UTR of the fragile X mental retardation 1 (*FMR1*) gene, which leads to its hypermethylation, transcriptional silencing, and subsequent lack of the fragile X mental retardation protein (FMRP) [5].

As a widely expressed RNA binding protein in the brain, FMRP plays a pivotal role in the stability and translational regulation of hundreds of mRNAs involved in synaptic protein synthesis [6], synaptic plasticity and neuronal development [7–9]. Absence or incorrect expression of FMRP is directly linked with alterations in dendritic spine architecture, synaptogenesis, and neural connectivity, as suggested by profound alterations of anatomical development in the cortex of *Fmr1* knock-out (*Fmr1* KO) mice [5,10]. In vivo studies on *Fmr1* KO mice have revealed many aspects of FXS pathophysiology, providing fundamental information on the functionality of FMRP and its involvement in neurogenesis, neuronal maturation and synaptic plasticity formation [11–13]. However, despite the *Fmr1* KO mouse represents a fundamental resource in understanding the molecular pathways altered in FXS, physiological and evolutionary species-specific differences have hampered translating these results from rodents to humans, raising the need for a humanized FXS model. In the last decade, human induced Pluripotent Stem Cells (hiPSCs) have emerged as useful tools for modeling neurodevelopmental disorders, including FXS [14–17]. Moreover, introducing specific disease-causing genomic alterations into the hiPSC line of choice by gene editing allows minimizing possible confounding effects due to the different genetic background of patient-derived hiPSC lines [18]. Notably, genome editing allows generating isogenic pools of wild-type and mutant hiPSC clones, providing a straightforward approach for unmasking phenotypic and functional alterations characteristic of complex diseases, such as Alzheimer’s disease [19], Parkinson’s disease [20], Huntington’s disease [21] and FXS [21].

While many neural cell types can be obtained by hiPSC differentiation, the lack of a three-dimensional (3D) tissue complexity fails to properly recapitulate the in vivo neurodevelopmental scenario in widely used bidimensional (2D) differentiation paradigms [22]. It has been shown that growing onto a flat and hard surface can alter gene expression and protein synthesis [23,24], drastically affecting crucial neurodevelopmental behaviors, such as differentiation and migration [25]. For this reason, although monolayer cultures are still widely used, novel 3D models have been developed to maintain more physiological conditions in terms of cellular morphology and cell-to-cell and cell-extracellular matrix contacts [26–28]. In this regard, by providing greater complexity and better fidelity to the environment in which neural cells reside in vivo, hiPSC-derived brain organoids may represent an important resource for the study of many neurodevelopmental diseases [29]. Brain organoids have been already used to model several neurodevelopmental disorders, including microcephaly [30], autism [31,32], Miller-Dieker syndrome [33], Timothy syndrome [34] and Angelman syndrome [35]. Nevertheless, despite the fast-growing use of this cutting-edge technology, to the best of our knowledge, the use of brain organoids to model FXS has not been yet pursued.

To investigate the function of FMRP in cortical brain development, we genetically engineered a healthy control hiPSC line by truncating the *FMR1* gene through CRISPR/Cas9 gene editing, thus generating an isogenic FMRP-KO hiPSC line. We report the phenotypical and functional characterization of FMRP-deficient hiPSC-derived cortical neurons, which display altered gene expression and impaired differentiation when compared with the isogenic control. Moreover, we generated and characterized 3D self-assembled brain organoids. Interestingly, since the brain organoid model recapitulates the early stages of human cortical development, all the phenotypical perturbations caused by FMRP deficiency, in terms of neuro-glia development, can be evaluated in a critical developmental period otherwise hardly accessible for investigations in humans. Moreover, the presence of intrinsically heterogeneous cell populations within brain organoids allowed assessing simultaneously the impact of FMRP loss on neuronal network and glial cells.

## MATERIALS AND METHODS

### Generation of the FMRP-KO hiPSC line

The pGEM-T donor vector (Promega) was engineered by introducing a donor construct, consisting of a cassette encoding for the self-cleavage peptide T2A, the puromycin resistance gene (PURO-resistance), and the beta-GH cleavage and polyadenylation site (pA). The cassette was amplified from AAVS1-Hb9-GFP vector [36] and flanked by FMR1 homology sequences to promote its integration in the genome upon homology-directed repair, interrupting the FMR1 coding sequence. A second vector, pX330-U6-Chimeric_BB-CBh-hSpCas9 (Addgene, ID #42230), coding for the nuclease Cas9 and the chimeric guide RNA (gRNA), was engineered inserting guide sequences targeting *FMR1* exon 2 (gRNA#1 Fw CACCGGTCCAGTTTGAGTGCTTTTC / Rv: AAACGAAAAGCACTCAAACTGGACC; gRNA#2 Fw: CACCGTTCCTGAAAAGCACTCAAAC / Rv: AAACGTTTGAGTGCTTTTCAGGAAC).

Cells were electroporated with 2 μg of donor construct and 3 μg of px330-Cas9 plasmid using a Neon Transfection System (Thermo Fisher Scientific) and the following settings: 1200 V, 30 ms, 1 pulse. After transfection, hiPSCs were cultured in presence of 10 μM ROCK inhibitor (Y27632; Enzo Life Science) for one day. Clonal selection was carried out with 0.5 μg/ml puromycin for 10 days, and surviving clones were individually passaged and characterized as described in Fig. 1A and 1B.

**Figure 1.**
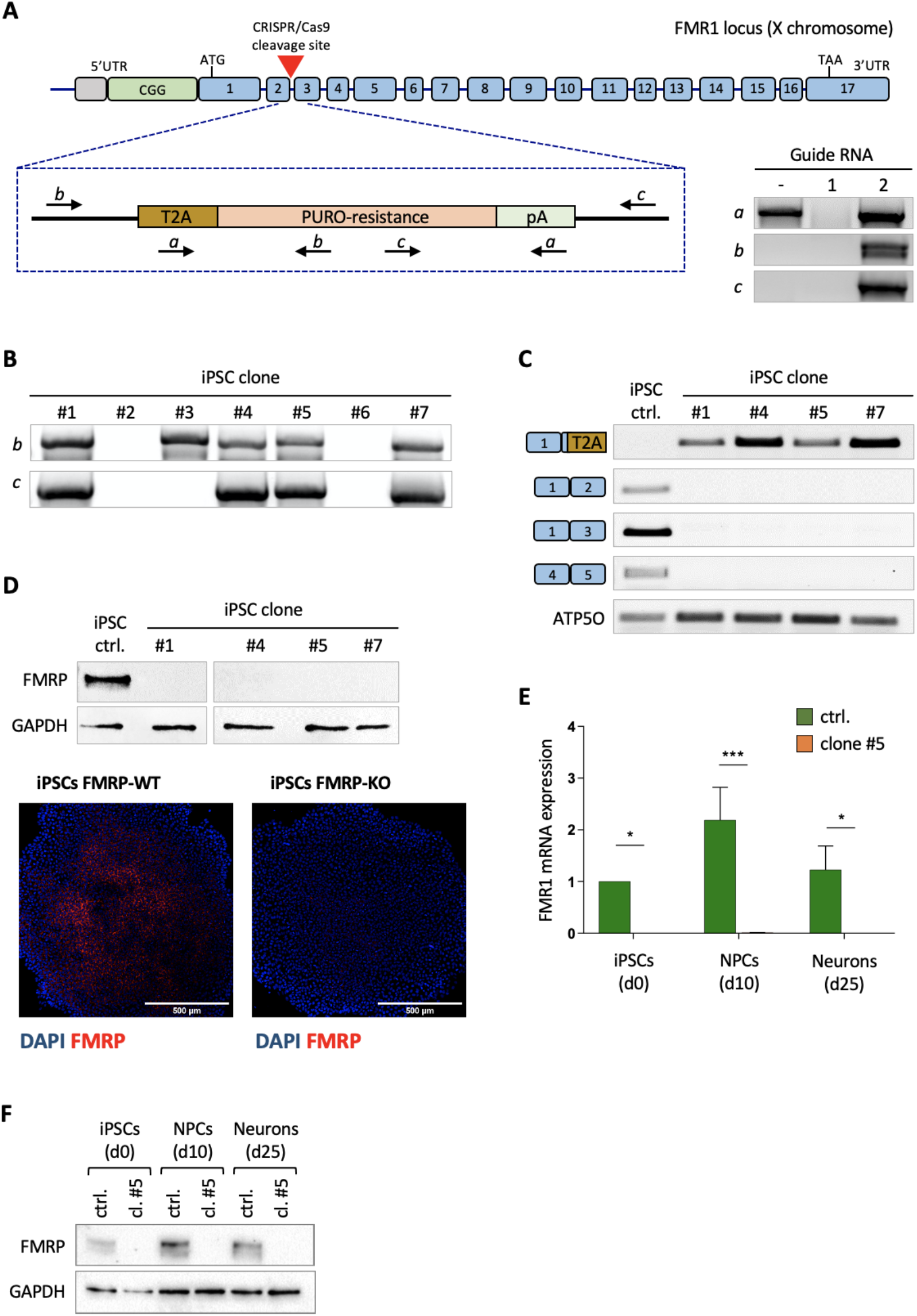
Generation and validation of FMRP knock-out hiPSCs. (A) Top: schematic representation of the human FMR1 locus and strategy for gene editing. The red triangle represents the site of cleavage by Cas9. The selection cassette, shown below, encodes for the self-cleavage peptide (T2A) and the puromycin resistance gene (PURO-resistance) and contains a cleavage and polyadenylation signal (pA). Primers used for the PCR analysis on genomic DNA are depicted as arrows. Bottom: PCR analysis on genomic DNA of hiPSCs transfected with the donor construct only (first lane) or co-transfected with the Cas9 and gRNA#1 or gRNA#2 (second and third lanes). (B) PCR analysis on genomic DNA of the indicated hiPSC clones with primers indicated in panel A. (C) RT-PCR analysis on RNA isolated from the indicated hiPSC clones with primers spanning exon-exon junctions. The housekeeping gene ATP5O was used as an internal control. (D) Top: western blot analysis of FMRP expression in the indicated hiPSC clones. GAPDH was used as loading control. Bottom: immunostaining analysis in parental and FMRP-KO hiPSCs. Red: FMRP; blue: DAPI. Scalebar: 500 μm. (E) Real-time qRT-PCR analysis of cortical neurons derived from parental and FMRP-KO hiPSCs (n=3 technical replicates; Student’s t test; paired; two tails; *p < 0.05, ***p < 0.001) (F) Western blot analysis of FMRP expression in cortical neurons derived from parental and FMRP-KO hiPSCs. GAPDH was used as loading control.

### Human iPSC maintenance and differentiation into cortical neurons

FMRP-WT and FMRP-KO hiPSC lines were cultured in Nutristem-XF (Biological Industries) supplemented with 0.1% Penicillin-Streptomycin (Thermo Fisher Scientific) onto hESC-qualified Matrigel (CORNING) functionalized plates. The culture medium was refreshed every day and cells were passaged every 4-5 days using 1 mg/mL Dispase II (Thermo Fisher Scientific).

hiPSCs were differentiated into cortical neurons according to a previously published protocol [37] with minor modifications. Briefly, hiPSCs were treated with Accutase (Thermo Fisher Scientific) and the single-cell suspension was plated in Matrigel-coated dishes at a seeding density of 100,000 cells per cm^2^, in Nutristem-XF supplemented with 10 μM ROCK inhibitor. The day after seeding the medium was changed to N2B27 medium consisting of DMEM-F12, Dulbecco’s Modified Eagle’s Medium/Nutrient Mixture F-12 Ham (Sigma Aldrich), Neurobasal Medium (Thermo Fisher Scientific), 1X N2 supplement (Thermo Fisher Scientific), 1X GlutaMAX (Thermo Fisher Scientific), 1X MEM-NEAA (Thermo Fisher Scientific), 1X B27 (Thermo Fisher Scientific), 1X Penicillin-Streptomycin (Thermo Fisher Scientific). This was considered as day 0 (D0). At this stage, SMAD inhibitors, 10 μM SB431542 and 500 nM LDN-193189 (Cayman Chemical), were added daily to induce neural fate. After 10 days, the uniform neuroepithelial sheet was broken into clumps of approximately 500 cells/clump and re-plated onto 1X poly-L-ornithine/laminin (Sigma Aldrich) coated dishes in N2B27 medium. At day 20, cells were dissociated using Accutase and plated onto poly-L-ornithine/laminin-coated dishes at a density of 100,000 per cm^2^ in N2B27 medium supplemented with 10 μM ROCK inhibitor for the first 24 hours. Culture medium was changed three times a week. At day 27, 2 μM Cyclopamine (Merck Life Science) was freshly added to N2B27 medium for 4 days. At day 30 to 35, immature neurons were re-plated into poly-L-ornithine/laminin-coated dishes at a density of 100,000 per cm^2^ in N2B27 medium supplemented with 20 ng/mL BDNF (Sigma Aldrich), 20 ng/mL GDNF (Peprotech), 200 ng/mL ascorbic acid (Sigma Aldrich), 1 mM cyclic AMP (Sigma Aldrich) and 5 μM DAPT (Adipogen Life Sciences).

### Generation of self-assembled brain organoids

A slightly modified version of Lancaster’s protocol [38] was used for the generation of cerebral organoids. After removing any spontaneously differentiated cells, hiPSC colonies were dissociated into a single-cell suspension and resuspended at a density of 180 cells/µL in Nutristem-XF with 10 μM ROCK inhibitor. 9,000 cells were then plated into each well of a low-attachment 96-well U-bottom plate (CORNING). After 48 hours, small embryoid bodies (EBs) of about 100-200 μm diameter were visible and the medium was switched to Neural Induction Medium (NIM) consisting of DMEM-F12 supplemented with 1X N2, 1X GlutaMAX supplement, 1X MEM-NEAA and 1X penicillin-streptomycin, 1 μg/mL of Heparin (Sigma Aldrich). EBs were fed every other day, being careful not to disrupt them, until optically translucent edges appeared confirming the neuroepithelium formation. On day 15, EBs were encapsulated into Geltrex (Thermo Fisher Scientific) droplets and each spheroid was positioned in a single well of a low-attachment 24 well plate (CORNING) in Differentiation Neural Medium (DMEM-F12, Neurobasal medium, 1X GlutaMAX supplement, 0.5X MEM-NEAA, 1X penicillin-streptomycin, 0.5X N2 and 0.5X B27 without vitamin A). After 4 days in static culture conditions, organoids were transferred to an orbital shaker (110 RPM), which facilitates the nutrient diffusion within the organoid, fed with the Maturation medium containing vitamin A (DMEM-F12, Neurobasal medium, 1X GlutaMAX supplement, 0.5X MEM-NEAA, 1X penicillin-streptomycin, 0.5X N2 and 0.5X B27 with vitamin A). The culture medium was changed every 3-4 days. During the whole growing period, the organoids were grown separately, each in a single well, to avoid aggregation.

### Brain organoids size analysis

Brain organoids images were collected in light transmission using an EVOS microscope (Thermo Fisher Scientific) equipped with a 4x objective. Fiji software was used for calculating the organoid surface area (expressed in mm^2^) and maximum diameter (expressed in mm).

### Immunostaining and image acquisition and analisys of 2D cultures

Monolayer cortical networks were fixed in 4% paraformaldehyde (PFA; Sigma Aldrich) solution for 15 minutes at room temperature and washed twice with 1X phosphate buffer solution (PBS; Thermo Fisher Scientific). Fixed cells were permeabilized with PBS containing 0.2% Triton X-100 (Sigma Aldrich) for 15 minutes and blocked with a solution containing 0.2% Triton X-100 and 5% Goat Serum (Sigma Aldrich) for 20 minutes at room temperature. Cells were then incubated overnight at 4°C with primary antibodies at the following dilutions: mouse anti-PAX6 (sc81649 Santa Cruz Biotechnology, 1:50), mouse anti-GFAP (MAB360 Merck Millipore, 1:500), chicken anti-MAP2 (ab5392 Abcam, 1:2000), rabbit anti-β-TUBULIN III (TUJ1) (T2200 Sigma Aldrich, 1:2000), rabbit anti-FMRP (4317 Cell Signaling, 1:50), mouse anti-VGLUT1 (135303 Synaptic Systems, 1:250), rabbit anti-PSD95 (3450 Cell Signaling, 1:250), mouse anti-GAD67 (sc-28376 Santa Cruz, 1:250). The day after, the primary antibody solution was washed out and cells were incubated with secondary antibodies for 1 hour at room temperature in the dark. AlexaFluor secondary antibodies (Thermo Fisher Scientific) were used at the concentration of 1:750 (Supplementary Table 1), and DAPI (Sigma Aldrich) was used for nuclei staining.

For synapses quantification of 2D cortical networks, fluorescence images were acquired on an Olympus iX73 microscope equipped with an X-Light V3 spinning disc head (Crest Optics), an LDI laser illuminator (89 North), an Evolve EMCCD camera (Photometrics) and a MetaMorph software (Molecular Devices). Images of 512×512 pixels (267×267 nm/pixel) were acquired with a 60x/NA 1.35 oil objective (Olympus) in stack with z-step of 0.5 μm. Quantification was performed on Fiji software using the “Puncta Analyzer” plugin. Stack images were flattened in a maximum intensity Z-projection and after removing the background noise, images were binarized allowing the count of synaptic puncta and the quantification of the co-localizing punctate.

For astrocytes quantification on 2D cortical populations, images were analyzed on Fiji software quantifying the GFAP signal as fluorescence intensity. The fluorescence threshold was adjusted to accurately represent the number of GFAP-positive cell processes. Integrated density, mean background value and the total area of the field of view (FOV) were calculated in order to get the mean fluorescence value per μm^2^.

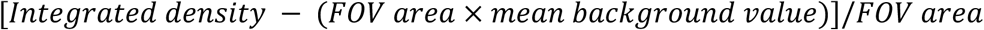

### Immunostaining and image acquisition and analisys of brain organoid

Brain organoids were collected at day 54 and 70, incubated in 4% PFA solution for 30 minutes and then moved in PBS with 30% sucrose overnight at 4° C. The following morning, the sucrose solution was removed and organoids were cryosectioned using a standard cryostat (Leica CM1860, Leica Biosystems). 50 μm thick sections were collected on Ultra Plus slides (Thermo Fisher Scientific) and stored at +4°C.

For immunostaining, organoid sections were quickly washed for 10 minutes in PBS and then incubated for 30 minutes with a warm antigen retrieval solution containing 1X Citrate buffer, at pH 6.0 (Sigma Aldrich). Samples were exposed to a 200 mM Glycine (Sigma Aldrich) solution in PBS for 15 minutes before permeabilization in 0.3% Triton-X in PBS for 45 minutes. After a double washing, sections were blocked in 0.3% Triton-X and 5% goat serum in PBS for 1 hour. Tissues were then incubated overnight with primary antibodies in 0.3% Triton-X and 5% goat serum in PBS at the following dilutions: mouse anti-PAX6 (sc81649 Santa Cruz Biotechnology, 1:50), rabbit anti-FOXG1 (ab18259 Abcam, 1:200), mouse anti-GFAP (MAB360 Merck Millipore, 1:500), chicken anti-MAP2 (ab5392 Abcam, 1:1000), rabbit anti-N-CADHERIN (ab18203 Abcam, 1:50), rabbit anti-TBR1 (20932-1-AP Proteintech, 1:150), rabbit anti-β-TUBULIN III (TUJ1) (T2200 Sigma Aldrich, 1:1000), rat anti-CTIP2 (ab18465 Abcam, 1:300). The day after, the primary antibody solution was washed out and cells were incubated with secondary antibodies for 2 hours at room temperature in a dark room. AlexaFluor secondary antibodies (Thermo Fisher Scientific) were used at the concentration of 1:750 (Supplementary Table 1), and Hoechst (Sigma Aldrich, 1:1000) was used for nuclei staining. Finally, after 3 washes with 0.15% Triton-X in PBS, organoid sections were mounted with ProLong Diamond Antifade Mountant (Thermo Fisher Scientific) and sealed with nail polish. The slides were stored at +4 ° C until acquisition.

Confocal images of GFAP, TUJ1 and MAP2 staining were acquired on Fluoview FV10i (Olympus) confocal laser scanning microscope. Multiarea images were acquired with a 10x/NA 0.4 phase contrast objective in stack with z-step of 10 μm. Each acquisition allowed to collect the signals present in the entire 3D slice of the organoid.

Images were analyzed on Fiji software quantifying the GFAP, MAP2 and TUJ1 signal as fluorescence intensity normalized for the area of the entire slice.

### Western blot

Whole-cell protein extracts were prepared from samples lysed in RIPA buffer (150 mM NaCl, 5 mM EDTA, 50 mM Tris Base, 1% NP-40, 0.5% Sodium Deoxycholate, 0.1% Sodium Dodecyl Sulfate). Extracts were separated by electrophoresis on NuPAGE 4-12% Bis-Tris gels (Thermo Fisher Scientific) in MOPS buffer (Thermo Fisher Scientific) and blotted onto nitrocellulose membrane (GE Healthcare). Immunoblots were incubated overnight with polyclonal rabbit anti-FMRP antibody (ab27455 Abcam, 1:1000) and for 1 hour with monoclonal mouse anti-GAPDH antibody (sc-32233 Santa Cruz, 1:3000) as loading control. Membranes were then incubated with HRP-conjugated donkey anti-mouse IgG (H+L) (IS20404 Immunological Sciences, 1:5000) or donkey anti-rabbit IgG (H+L) (IS20405 Immunological Sciences, 1:5000) secondary antibodies for 45 minutes. HRP signal was revealed with the Clarity Western ECL Substrates kit (Bio-rad). Images were acquired with the Chemidoc MP (Bio-Rad) and analyzed with the ImageLab software (Bio-Rad).

### PCR, RT-PCR and RT-qPCR

Genomic DNA was extracted from cells using PCRBIO Rapid Extract PCR Kit (PCR Biosystems) according to manufacturer instructions. After extraction, genomic DNA was amplified for 40 cycles using oligos indicated in Supplementary Table 2.

Total RNA was extracted with the EZNA Total RNA Kit I (Omega Bio-Tek) and retrotranscribed using the iScript Reverse Transcription Supermix for RT-qPCR (Bio-Rad).

For FMRP-KO clones screening (Fig. 1C), target cDNA sequences were amplified for 35 reaction cycles with the enzyme MyTaq DNA Polymerase (Bioline). The cDNA of the housekeeping gene *ATP5O* (ATP synthase, H+ transporting, mitochondrial F1 complex, O subunit), used as internal control, was amplified for 28 reaction cycles.

Real-time RT-PCR was performed with iTaq Universal SYBR Green Supermix (Bio-Rad) on a ViiA 7 Real-Time PCR System (Applied Biosystems). A complete list of primers is provided in supplementary material (Supplementary Table 2).

### Calcium imaging recordings and data processing

Fluorescence images were acquired at room temperature using a customized digital imaging microscope. Excitation of calcium dye was achieved at wavelenght 488 nm using the highly stable light source Lambda XL (Sutter Instrument) equipped with a Lambda 10-B optical filter changer (Sutter Instrument). For the collection of the emitted light, a 525/50 nm filter was used. Fluorescence was visualized using the Zeiss Axio observer A1 inverted microscope (Zeiss) equipped with a Zeiss A-Plan 10x/NA 0.25 infinity corrected objective (Zeiss) and a CoolSNAPHQ2 camera (Photometrics). Images acquisition at 4 Hz sampling rate were performed using Micromanager software. Changes in the intracellular Ca^2+^ level were observed using the high-affinity Ca^2+^-sensitive indicator Fluo4-AM (Invitrogen) which was used at the concentration of 5 μM by incubating neuronal cultures for 30 minutes at 37°C in HEPES-buffered external solution (NES) containing 140 mM NaCl, 2.8 mM KCl, 2mM CaCl_2_, 2mM MgCl_2_, 10mM HEPES, 10 mM D-glucose (pH 7.3 with NaOH; 290 mOsm).

Calcium imaging data processing was performed through custom numeric codes implemented in MATLAB environment. Recorded images sequences were collected and saved as 3D matrix. The first step of analysis concerned the definition of regions of interest for neurons. We realized an automated algorithm to identify only active neurons by analyzing the cumulative difference of the signal between the various frames of the time series. The obtained matrix was studied in the frequency domain via 2D Fourier transform and suitably filtered to eliminate high-frequency noisy components; then the local maxima of the matrix were selected. The algorithm provided the possibility to manually adjust eventual false or missing detections. Once established active cell positions (no more than 200 cells selected for each field of view), their fluorescence signals as a function of time were collected. A supervised analysis allowed the detection of calcium transients and relative characteristics (amplitude, rising and decay time). In detail, the raw traces of the neurons extracted from the t-stack were baseline corrected and normalized as ΔF/F_0_; then smoothed using a moving average filter. On the filtered traces, a putative event was detected when a series of conditions were satisfied: at the onset the fluorescence intensity and the slope of the trace show an increase; at the offset the slope of the trace decreases and a certain time interval occurs within the onset and the offset [39]. Threshold for peaks amplitude was set to 1% of the baseline value. This operation gave a first indication about the onset time of each event, which was used as starting point for a fitting procedure based on a previously described algorithm [40]. The algorithm assumes that there exists an elementary calcium transient characteristic of a single action potential and that the calcium transients add up linearly. Each putative transient was fitted in two-step with a model function composed of a single-exponential rise and a single-exponential decay.

In the first step, we fitted the onset to determine the start of the event t_0_ and the onset time constant τ_on_.

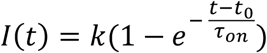

Then we fitted the entire calcium transient to estimate amplitude and the decay component τ_off_.

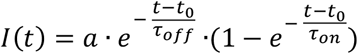

The analysis of calcium dynamics was crucial to distinguish fast calcium transients, typical of neuronal activity, from calcium signals characterized by a slowest onset (release from internal stores, calcium signals from other cell types). According to our experimental conditions, we established a threshold τ^*^=2 s to recognize and discard non-neuronal signals: neurons whose mean rising time results below this threshold were selected, all other signals were not considered for final statistics. Event rate, amplitude and synchrony of the network (evaluated as the relative number of simultaneous events) were exported in Microsoft Excel.

For the agonist-evoked calcium activity, images were acquired at a frequency of 1 Hz for 15 minutes with an exposure time of 200 ms through a BX51WI microscope (Olympus Corporation, Tokyo, JP. Objectives: LUMPlanF N 10x/0.10, air, and 40x/0.80, water immersion, Olympus Corporation). Fluo4-AM was excited at 488 nm with an Optoscan monochromator (Cairn Research, Facersham, UK). Light was generated by a xenon lamp Optosource (Cairn Research). A borosilicated glass micropipette was filled with NES and 2 mM Glutamate or 1 mM GABA (Sigma Aldrich), and moved via a micromanipulator MP-225 (Sutter Instruments, Novato, CA) to reach the core of the field of view, around 50 µm beneath the surface of the dish.

The basal fluorescence was evaluated for 5 minutes, then a small volume of agonist-containing solution was puffed on the cells via a pneumatic pico-pump (PV820; World Precision Instruments, Inc., Sarasota, FL) with a brief pressure (10 psi; 100 ms). The images were acquired with a camera CCD CoolSnap MYO (Photometrics, Tucson, AZ, USA) and then analyzed with MetaFluor software as fluorescence variation measured into regions of interest (ROI) corresponding to single cells. To quantify the signal, the formula (F-F_0_)/F_0_ was used, where F_0_ is the average fluorescence before the agonist application and F the fluorescence during the time-lapse acquisition.

### Statistical data analysis

Statistical analysis, graphs and plots were generated using GraphPad Prism 6 (GraphPad Software) and MATLAB 2016b (MathWorks). To verify whether our data sets were reflecting normal distribution, the Shapiro-Wilk normality test was performed. Where the normality distribution was not fulfilled, statistical significance analysis was performed using the nonparametric two-sided Mann–Whitney test (MW test, P=0.05). In all other cases, wheter not stated otherwise, t-Student test (P=0.05) was performed, and data set are given as mean ± standard error of the mean (s.e.m.).

Statistical significance of the cumulative distribution were calculated using the nonparametric Kolmogorov-Smirnov test.

## RESULTS

### Generation and characterization of FMRP KO hiPSCs

In order to produce an in vitro FXS model system made of isogenic mutant and control hiPSC lines, we generated a FMRP knock-out line by gene editing in a FMRP wild-type genetic background. The human *FMR1* genomic locus is depicted in Figure 1A. We designed a strategy, based on the CRISPR/Cas9 system, to block the production of FMRP at the level of transcription. Taking into consideration the presence of an antisense non-coding RNA, the production of multiple transcripts from the *FMR1* gene and the apparent absence of downstream alternative transcription start sites (Supplementary Figure S1), the strategy to produce a knock-out mutant was based on the insertion of a selection gene spliced in frame with the first *FMR1* exon. We designed two guide RNAs (gRNA#1 and gRNA#2) directed against this region and a donor construct with homology arms. Upon homology-directed repair, a selection cassette encoding for a self-cleavage peptide (T2A), a puromycin resistance gene and a cleavage and polyadenylation signal was inserted in frame after the first three codons of exon 2. With this design, as a result of transcription and splicing, a transcript encoding for the puromycin resistance protein (puromycin-N-acetyltransferase) would be produced, while no FMRP mRNA could be generated.

The fact that the *FMR1* gene is on the X-chromosome facilitated the knock-out strategy in male cells (the hiPSC line WTI [41]), as only one allele needed to be edited. After transfection and selection with puromycin, PCR analysis on genomic DNA with specific primers showed effective integration of the cassette in hiPSCs co-transfected with the Cas9, the donor construct and gRNA#2 (Figure 1A). We next isolated single clones from the gRNA#2-transfected population. As shown in Figure 1B, PCR analysis on genomic DNA confirmed the specific integration of the cassette in several clones. We then performed on these clones a RT-PCR analysis with primer pairs flanking splicing junctions (Figure 1C). All clones showed the amplification product of the primer set specific for the spliced knock-out construct (first row in Figure 1C). Conversely, the unedited spliced mRNA could be detected in the parental line only, as demonstrated by the analysis with reverse primers annealing beyond exon 1 (Figure 1C, second to fourth row). Consistently, in the same cells no FMRP protein was detectable by western blot or immunostaining (Figure 1D). Complete absence of *FMR1* transcript and protein was then confirmed in cortical neurons generated by gene edited hiPSC differentiation at day 10 and 25 (Figure 1E, 1F and 2A).

**Figure 2.**
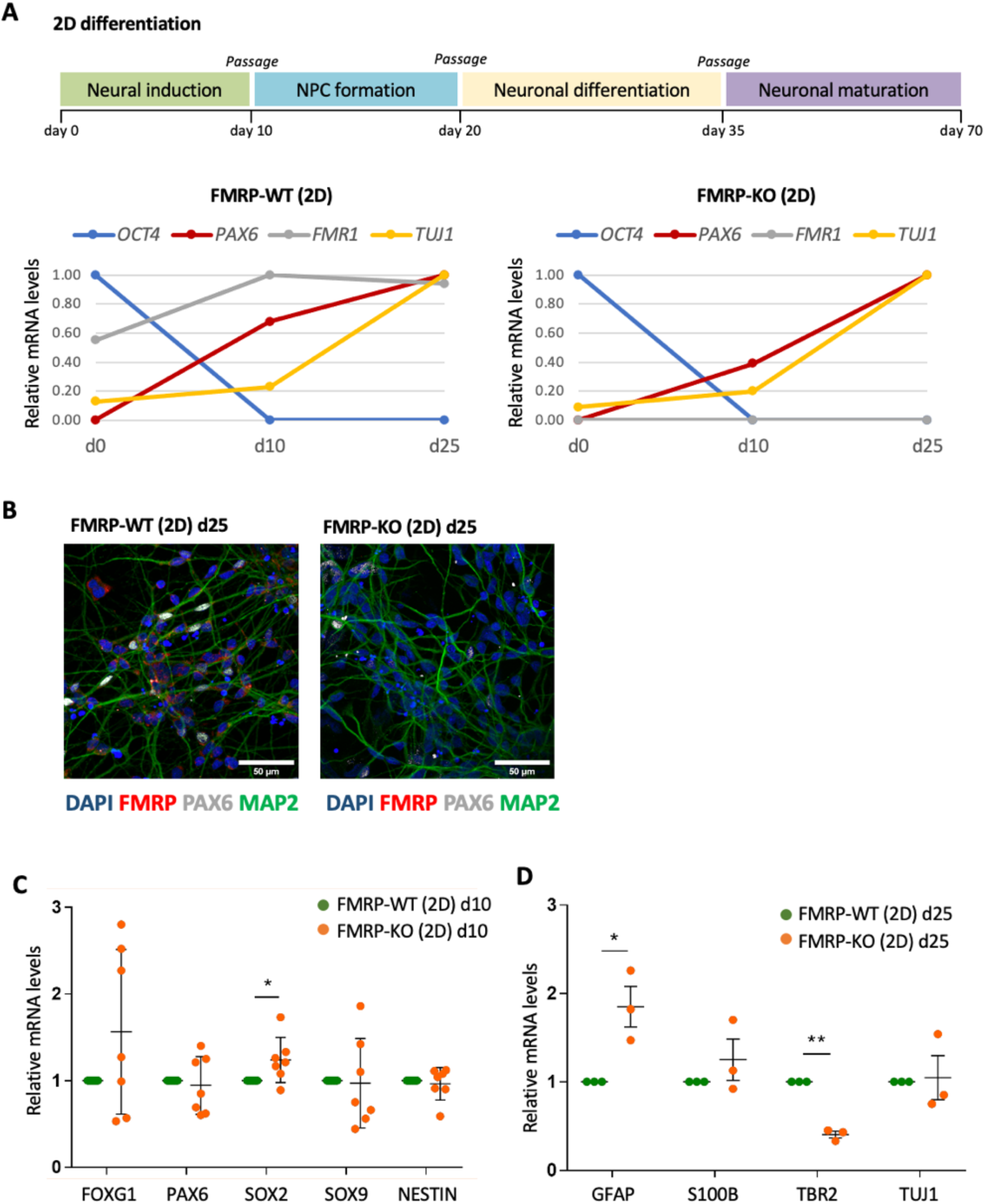
Differentiation of FMRP knock-out and isogenic control hiPSCs into cortical neurons. (A) Top: schematic representation of the cortical neuron differentiation protocol. Bottom: real-time qRT-PCR analysis of the indicated markers expression in differentiating FMRP-WT and FMRP-KO cells at the indicated time points. (B) Immunostaining analysis of PAX6 and FMRP expression in FMRP-WT and FMRP-KO cells at day 25. Scalebar: 50 µm. (C,D) Real-time qRT-PCR analysis of the indicated markers expression in FMRP-WT and FMRP-KO cells at day 10 (C) and 25 (D) of differentiation (each point represents a differentiation experiment; Student’s t test; paired; two tails; *p < 0.05, **p < 0.01).

One gene edited clone (clone #5; hereafter FMRP-KO) and parental isogenic hiPSCs (hereafter FMRP-WT) were differentiated to cortical neurons using a conventional protocol and 2D culture conditions. Acquisition of a neural cortical fate was induced by dual SMAD inhibition followed by the block of Hedgehog signaling with cyclopamine (Figure 2A, top) [37,42]. Exit from pluripotency and acquisition of neuronal character was assessed by qRT-PCR analysis of pluripotency (*OCT4*), neural progenitor cells (*PAX6*), and neuronal (*TUJ1*) markers (Figure 2A, bottom). Immunostaining analysis confirmed expression of PAX6 at the neural rosette stage and lack of FMRP in the FMRP-KO line (Figure 2B). We then analyzed a panel of early and late neural markers over multiple differentiation experiments to assess whether the absence of FMRP could lead to defects in differentiation at this stage. At day 10 of differentiation, corresponding to the late neural induction phase and initial acquisition of the neural progenitor cell character, we noticed a trend of increased expression for *SOX2* and *FOXG1* in FMRP-KO cells, with a substantial degree of variability among different biological replicates (Figure 2C). However, no significant change was observed for other early neural markers (*PAX6, SOX9* and *NESTIN*). At day 25, corresponding to a stage in which differentiating cells that are acquiring a neuronal character exit from the cell cycle, we noticed a significant increase in the levels of the astrocyte marker *GFAP* and concomitant decrease of the neuronal precursor marker *TBR2* in FMRP-KO cells, which also showed a slight but not significant increase of another astrocyte marker, *S100B* (Figure 2D). Taken together, these results validate a novel FMRP knock-out hiPSC line, generated by gene editing, showing early imbalance of neuronal and astrocyte markers during cortical differentiation.

### FMRP-KO cortical neurons display increased excitatory transmission during in vitro maturation

As cortical hyperexcitability is one of the hallmark of FXS, and neuronal hyperexcitability has been reported both in slices and culture of *Fmr1* KO mouse model [8], we characterized, through confocal analysis of immunofluorescence signals and functional recordings of calcium dynamics, the development of the glutamatergic and the GABAergic systems in 2D cortical cultures differentiated from FMRP-WT and FMRP-KO hiPSCs.

At a late stage of development and maturation a mixed population of differentiating cells was achieved, with the simultaneous presence of neural progenitors, neurons and glial cells. In both FMRP-WT and KO cultures at day 54, consistent with previous findings [37], immunofluorescence analysis of neuronal and glial markers indicated that GFAP staining was barely detectable (not shown), while the large majority of cells were positive for the neuronal marker MAP2 (Figure 3A, 3B). The presence of glutamatergic and GABAergic neurons was demonstrated by the positive staining of neurons for both presynaptic VGLUT1 and GAD67, respectively. Moreover, positive staining for PSD95 demonstrated the presence of post-synaptic glutamatergic specialization. When comparing FMRP-WT and FMRP-KO cultures at day 54, while we did not find any difference in presynaptic GAD67 staining (Figure 3A, 3B), we observed that FMRP-KO neurons display higher number of pre- (VGLUT1) and post- (PSD95) synaptic glutamatergic components (puncta), with increased colocalization compared to WT, indicating faster excitatory synaptic development (Figure 3A, 3B), thus indicating that during cortical neuron maturation FMRP-KO neurons display an accelerated development of glutamatergic system.

**Figure 3.**
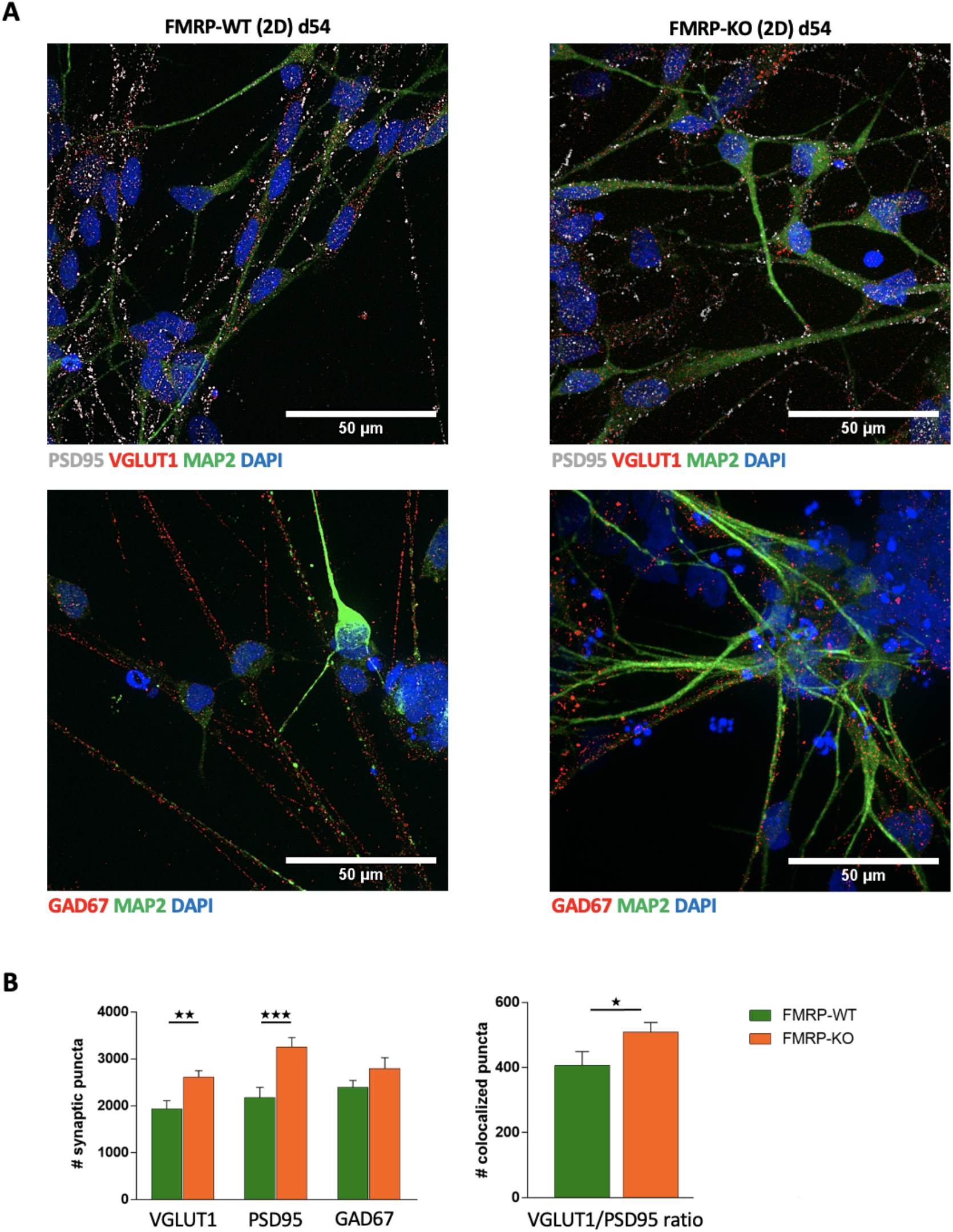
Immunofluorescence analysis of 2D cortical cultures at day 54. (A) Representative immunofluorescence images of glutamatergic and GABAergic synapses at day 54 in 2D cortical cultures differentiated from FMRP-WT and FMRP-KO hiPSC. Top: glutamatergic synaptic puncta identified as positive for VGLUT1 (pre-synaptic, red) and PSD95 (post-synaptic, white) respectively. Bottom: GABAergic pre-synaptic puncta, positive for GAD67 (red). Nuclei were stained with DAPI (blue) and the neurite branches with MAP2 (green). Scale bar: 50 µm. (B) Bar charts representing (left) the quantification of glutamatergic and GABAergic synaptic puncta (1929 ± 180 VGLUT1 puncta in WT and 2610 ± 139 VGLUT1 puncta in KO, **p<0.01 KO vs WT, MW-test; 2168 ± 223 PSD95 puncta in WT and 3255 ±200 PSD95 puncta in KO, ***p<0,001 KO vs WT; 2394 ± 145 GAD67 puncta in WT and 2792 ± 234 GAD67 puncta in KO, not significant; n=20 FOVs for each condition) and (right) VGLUT1/PSD95 colocalization ratio as in (A), in both genotypes (407 ± 41 VGLUT1/PSD95 colocalized puncta in WT and 509 ± 29 VGLUT1/PSD95 colocalized puncta in KO, *p<0.05 KO vs WT; n=20 FOVs for each condition).

At day 70, side-by-side comparison of FMRP-WT and FMRP-KO indicated another early FXS phenotype, consisting of an accelerated specification of GFAP-positive astrocytes as revealed by GFAP staining (Figure 4A, 4B), while the expression levels of pre- (VGLUT1) and post- (PSD95) synaptic glutamatergic markers became similar, suggesting that the increase in glutamatergic network was transient in FMRP-KO cultures (Figure 4C, 4D).

**Figure 4.**
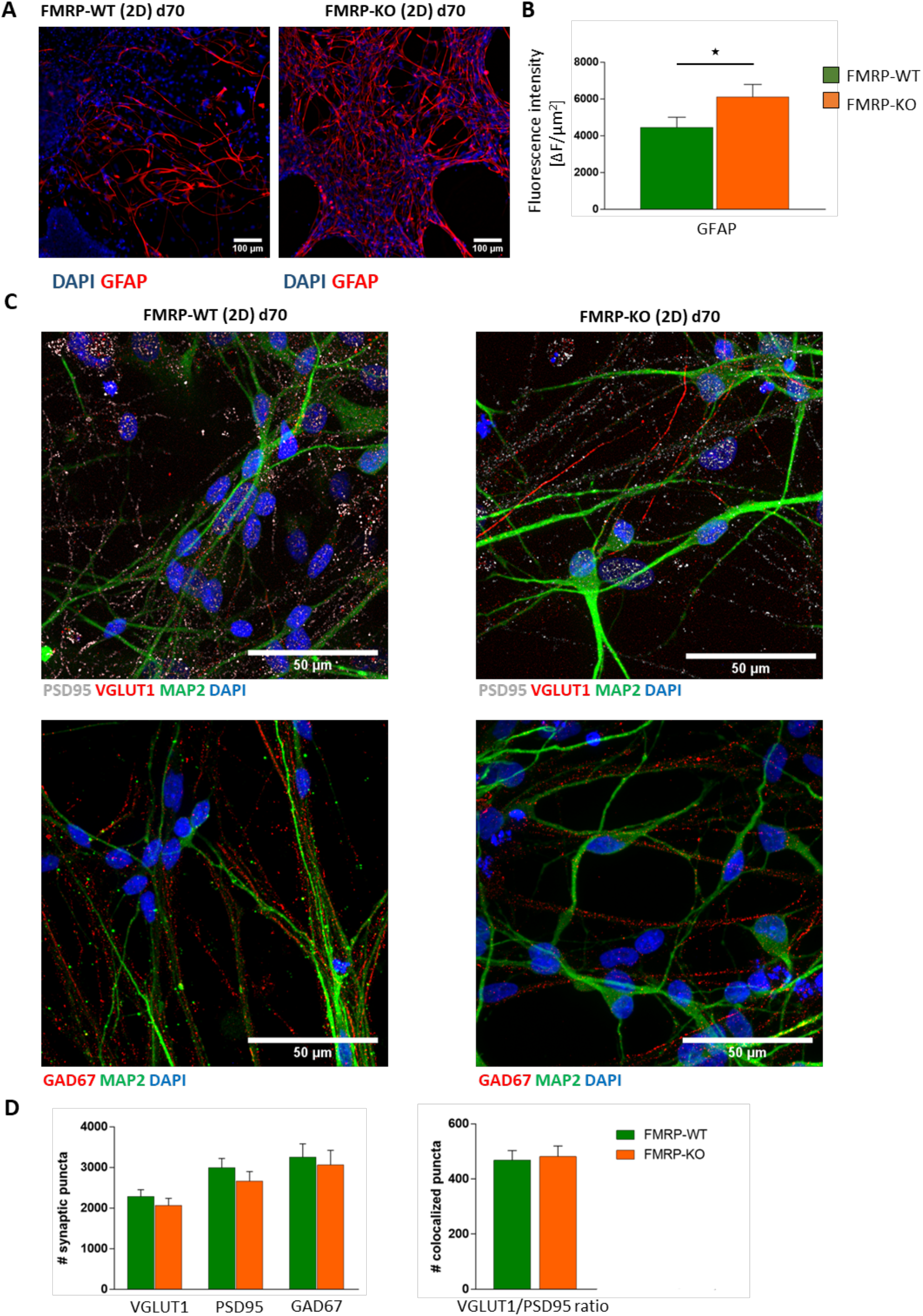
Immunofluorescence analysis of 2D cortical cultures at day 70. (A) Representative images of differentiated 2D cortical cultures from FMRP-WT and FMRP-KO hiPSCs immunolabeled with anti-GFAP antibody (red) and DAPI for nuclei visualization (blue). Scale bar: 100 μm. (B) Quantification of GFAP area covered by fluorescent signal/field of view (*p<0.05 KO vs WT, MW-test; n = 20 FOV). (C) Representative images of glutamatergic and GABAergic synapses at day 70. Top: glutamatergic synaptic puncta identified as positive for VGLUT1 (pre-synaptic, red) and PSD95 (post-synaptic, white) respectively. Bottom: GABAergic pre-synaptic puncta, positive for GAD67 (red). Nuclei were stained with DAPI (blue) and the neurite branches with MAP2 (green). Scale bar: 50 µm. (D) Bar charts representing (left) the quantification of glutamatergic and GABAergic synaptic puncta (2277 ± 176 VGLUT1 puncta in WT and 2066 ± 172 VGLUT1 puncta in KO, not significant; 2996 ± 224 PSD95 puncta in WT and 2665 ±234 PSD95 puncta in KO, not significant; 3248 ± 334 GAD67 puncta in WT and 3059 ± 361 GAD67 puncta in KO, not significant; n=20 FOVs for each condition) and (right) VGLUT1/PSD95 colocalization ratio as in (A), in both genotypes ((467 ± 35 VGLUT1/PSD95 colocalized puncta in WT and 481 ± 38 VGLUT1/PSD95 colocalized puncta in KO, not significant; n=20 FOVs for each condition).

Strikingly, at day 70, we observed a functional FXS phenotype when measuring spontaneous intracellular calcium dynamics at network level and the glutamate and GABA evoked calcium responses at the level of single neuron. We recorded spontaneous calcium transients in the 2D neuronal networks loaded with Fluo-4M, using a customized digital fluorescence microscope with a 4 Hz sampling of a large field of view (260×200 micron), that allowed to monitor a large number of cells at the same time. Data from a 5 minute time-lapse recordings were collected and analyzed through a custom-made algorithm (Supplementary Figure S2) designed to recognize cells, select active cells, sort calcium events, dividing cells in two groups characterized by the occurrence of calcium events with fast or slow rise time respectively (Figure 5A, 5B), and finally extract functional properties such as amplitude, frequency, kinetic parameters and network synchronicity [42,43].

**Figure 5.**
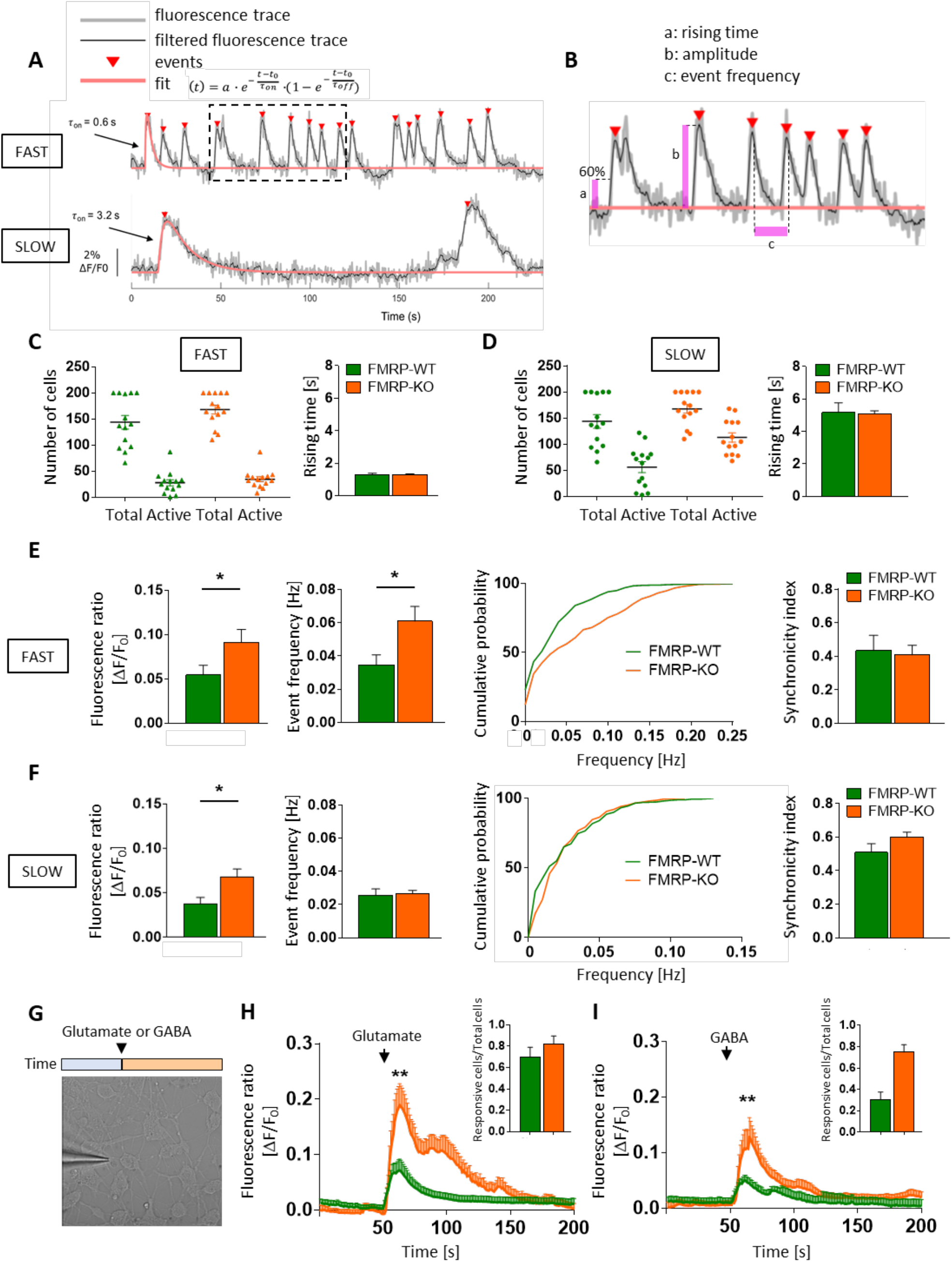
Spontaneous and evoked calcium activity analysis of 2D cortical cultures at day 70. (A) Calcium traces as a function of ΔF/F_0_ of FAST (top) and SLOW (bottom) active cells. (B) Representative calcium trace (insert from panel A) showing extracted functional properties: rising time (a), amplitude (b) and event frequency (c). (C) FAST active cells with respect to total cells in both FMRP-KO and WT populations (n=14 FOVs for genotype; 144 ± 13 total cells in WT, 28 ± 6 active cells in WT; 168 ± 8 total cells in KO, 34 ± 5 active cells in KO; rising time 1,3 ± 0,1 sec in WT and 1,3 ± 0,03 sec in KO). (D) SLOW active cells with respect to total cells in both FMRP-KO and WT populations (n=14 FOVs for genotype; 144 ± 13 total cells in WT, 56 ± 11 active cells in WT; 168 ± 8 total cells in KO, 113 ± 9 active cells in KO; rising time 5,2 ± 0,6 sec in WT and 5,1 ± 0,17 sec in KO). (E) Graph plots reporting the functional properties of FAST active cells: amplitude (left, n=14 FOVs, *p<0,05 KO vs WT, MW test), frequency and relative cumulative distribution (middle, n=14 FOVs, 0,035 Hz in WT and 0,06 Hz in KO, *p<0,05 KO vs WT) and synchronicity (right) of spontaneous calcium events of FMRP-WT and FMRP-KO fast cells (n=14, 0,43 ± 0.09 in WT and 0,41 ± 0.06 in KO not significant). (F) Graph plots reporting the functional properties of SLOW active cells amplitude (left, n=14 FOVs, *p<0,05 KO vs WT, MW test), frequency and relative cumulative distribution (middle, n=14 FOVs, 0,025 Hz in WT and 0,025 Hz in KO, not significant) and synchronicity (right) of spontaneous calcium events of FMRP-WT and FMRP-KO slow cells (n=14, 0,51 ± 0.05 in WT and 0,6 ± 0.03 in KO, not significant). (G) Representative FOV showing a puffer pipette for local Glutamate and GABA application with a cartoon showing the experimental approach. (H) Glutamatergic calcium transient response of FMRP-KO and WT cortical neurons after puffer applications (n=5 FOVs, 23 ± 4,7 total cells and 17 ± 4 active cells in WT; n = 4 FOVs, 22 ± 2 total cells and 18 ± 1 active cells in KO; **p<0.01 KO vs WT). (I) GABAergic calcium transient response of FMRP-KO and WT cortical neurons after puffer applications (n=5 FOVs, 22,6 ± 3,5 total cells and 7 ± 1,8 active cells in WT; n=5 FOVs, 14,6 ± 1,7 total cells and 11,4 ± 2 active cells in KO, **p<0.01 KO vs WT).

For each field of view (see Supplementary video 1 and 2), using 2 seconds value as rising time threshold, we divided active cells in FAST (Figure 5A top) and SLOW (Figure 5A bottom) population. As represented in the dot plots, the proportion of both FAST and SLOW active cells were similar in FMRP-WT and FMRP-KO cultures (Figure 5C, 5D). As reported in Figure 5E the spontaneous activity of FAST cells, most probably neurons, was significantly stronger in FXS cultures; indeed both the amplitude (Figure 5E left) and the frequency (Figure 5E middle and right) of calcium events were significantly higher in FMRP-KO neurons respect to FMRP-WT neurons, with no difference in synchronized firing, thus suggesting the possible overexcitability of the FXS cortical network. On the other hand, analyzing calcium traces extracted from the SLOW population, likely ascribable to glial cells and differentiating neuronal cells, we found that FMRP-KO cells displayed spontaneous calcium events characterized by higher amplitude (Figure 5F left) but similar frequency (Figure 5F middle and right) and synchronization with respect to FMRP-WT cells, suggesting that also the astrocytic network is more active in FXS cortical cultures.

As the cortical network activity and excitability are strictly dependent on the balance of excitatory and inhibitory transmission, we evaluated the potency and the sign of glutamatergic and gabaergic transmission monitoring in 2D cultures calcium transients evoked by a local and brief agonist application through a puffer pipette (Figure 5G). Glutamate application (2 mM, 100 ms, 10 p.s.i.) evoked intracellular calcium responses, indicating functional expression of glutamate receptors (Figure 5H, insert). As depicted in Figure 5H glutamatergic responses were significantly higher in FMRP-KO cells with respect to FMRP-WT, suggesting that even though the number of pre- and post-synaptic markers was similar in the two genotypes, the neuronal depolarization mediated by glutamatergic receptor activation is strongly upregulated in our FXS model, probably contributing to the increased network activity. Neuronal hyperexcitability and seizure susceptibility are common in FXS patients and animal models, and they are believed to result from an imbalance between excitatory and inhibitory drives in intracortical circuits, with reduced inhibition as key mechanism for circuit hyperexcitability. As defective GABAergic inhibition may arise from a depolarizing GABA effect, and depolarizing GABA is typical of immature neurons, we tested if GABA application may induce a transient calcium increase in our cultures at day 70. As reported in Figure 5I calcium transients evoked by puffer application of GABA (1 mM; 100 ms 10 psi) were significantly higher in FMRP-KO neurons with respect to FMRP-WT; moreover, while GABA application induced a depolarizing effect on 30±3% of FMRP-WT neurons (n=33 cells/ 6 FOV; Figure 5I, insert), in FMRP-KO cultures 75 ±3% (n=57 cells/ 6 FOV; Figure 5I, insert) of neurons responded to GABA with a depolarization mediated calcium transient, indicating that FMRP-KO cultures displayed an immature phenotype.

Altogether these data suggest that FMRP-KO cultures are characterized by a FXS-related functional phenotype with pronounced hyperexcitability mediated by increased glutamatergic transmission and decreased GABAergic inhibition.

### Development of a brain organoid-based FXS 3D disease model

The next step was to improve the hiPSC-based FXS model moving to a more complex, physiologically relevant in vitro model of the disease such as brain organoids.

Figure 6A shows the timeline of the experimental strategy used to generate self-assembled brain organoids from FMRP-WT and FMRP-KO hiPSCs. After 70 days in culture, brain organoids displayed typical laminar structures, as represented in Figure 6B. During organoids maturation, at day 50 we observed the appearance of markers typical of the ventricular and subventricular zones as N-Cadherin, FOXG1 and PAX6 (Figure 6C) confirming the presence of a germinal layer; moreover, developing cortical plates were positive for deep layer cortical neurons as revealed by the presence of CTIP2 and TBR1 positive cells, suggesting the initial formation of the cortical stratification. At day 70, the positive staining for TUJ1, MAP2 and GFAP revealed the presence of neuronal and glial cells, validating our protocol for cortical organoids generation.

**Figure 6.**
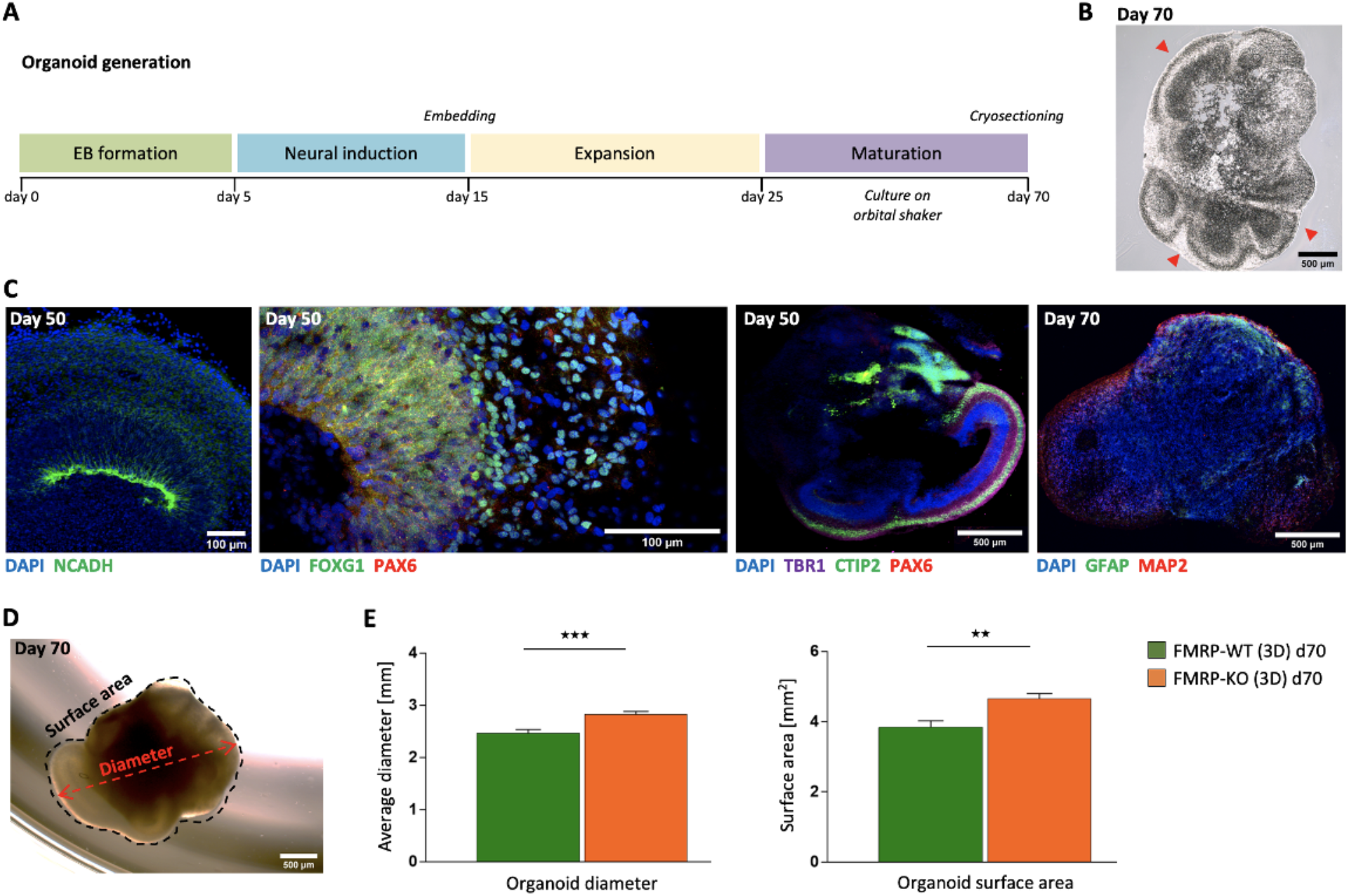
Evaluation of cellular identity and morphology in 3D brain organoid. (A) Scheme illustrating the timeline of development used for the brain organoid generation. (B) Representative image of day 70 cryosectioned FMRP-KO brain organoid. Arrows indicate the presence of developing cortical plates within the organoid. (C) Representative immunostaining highlighting the presence of a germinal layer positive for N-Cadherin and PAX6 markers (left and middle-left; scale bar: 100 μm) and an early-born deep-layer cortical layer positive for TBR1 and CTIP2 markers (middle-right and right; scale bar: 500 μm). Nuclei were stained with DAPI (blue). (D) Representative tracing of organoid maximum diameter and surface area at day 70. (E) Quantification of average diameter (n=40, d = 2,5 ± 0,07 mm in WT and n=77, d = 2,8 ± 0,06 mm in KO, ***p<0,001 KO vs WT) and average surface area (n=40, A = 3,8 ± 0,2 mm^2^ in WT and n=77, 4,7 ± 0,15 mm^2^ in KO, **p<0,01 KO vs WT) at day 70.

As the FMRP deficiency is believed to alter the early developmental stages of the corticogenesis, we compared cortical organoids obtained from FMRP-WT and FMRP-KO iPSCs and observed significantly higher diameter and surface area in FMRP-KO organoids at day 70 (Figure 6D, 6E).

Moreover, the characterization of the cellular identity within the cortical plates (Figure 7A), indicated that while the staining for TUJ1 was similar in the two genotypes (Figure 7B, left), FMRP-KO cerebral organoids displayed a reduced MAP2 staining (Figure 7B, middle) and increased GFAP labeling (Figure 7B left), reminiscent to what observed in 2D cultures (Figure 2D, 4A). At day 70 CTIP2 positive staining revealed the presence of layer V cortical neurons in both FMRP-WT and KO organoids (Supplementary Figure 3).

**Figure 7.**
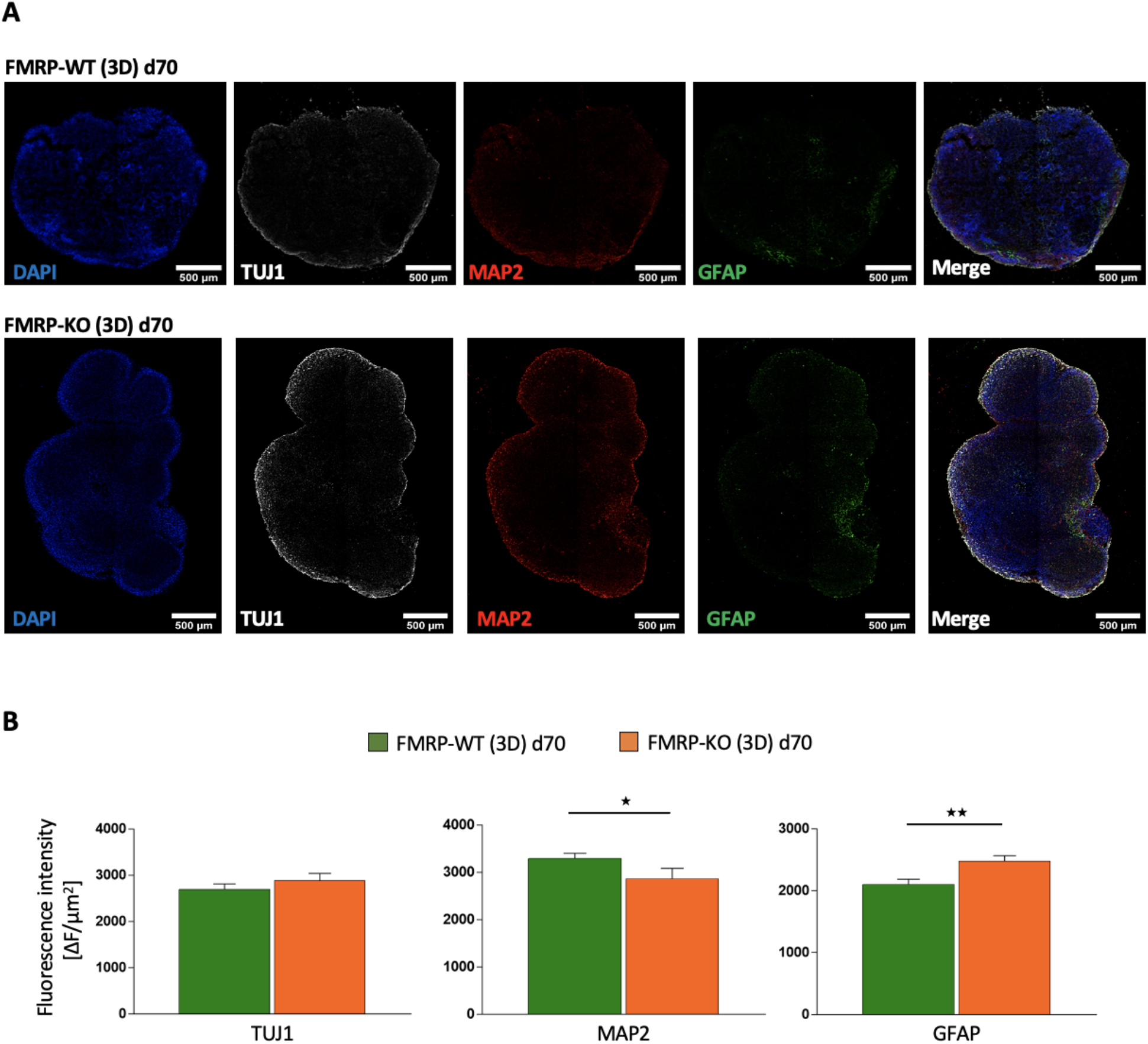
Immunofluorescence analysis of 3D brain organoids at day 70. (A) Cellular identity characterization in FMRP-WT and FMRP-KO brain organoids. Immunostaining at day 70 revealed cells positive for both pan-neuronal markers TUJ1 (white) and MAP2 (red), and also the presence of a GFAP (green) expressing astrocytes. Nuclei were stained with DAPI (blue). Scale bar: 500 μm. (B) Fluorescence intensity quantification of TUJ1, MAP2 and GFAP signals (n=20 slices/ 3 FMRP-WT organoids; 18 slices/ 4 FMRP-KO organoids; for MAP2 signal *p<0,05 KO vs WT, MW test; for GFAP signal, **p<0,01 KO vs WT).

These data suggest that a FXS phenotype can be obtained in 3D self-assembled brain organoids that, with respect to isogenic parental WT line, display bigger size and accelerated maturation of GFAP-positive astrocytes and reduced expression of the dendritic marker MAP2.

## DISCUSSION

The development of effective drugs for FXS is challenging because of the limited understanding of its pathophysiology, the differences between patients and animal models, and the difficulties in modelling FXS in vitro and in vivo. Progresses in disease-relevant hiPSC generation and modification by genome editing and in the generation of 3D organoids provide novel possibilities for disease modelling and drug screening for FXS. This study reports the generation of 2D and 3D in vitro FXS models from control and isogenic FMRP-KO hiPSCs, which recapitulate in vitro relevant FXS phenotypes. The long term aim is to overcome the issues due to differences between human patients and animal models. Indeed, a proper in vitro human disease model in which FXS evidences could be confirmed and implemented is still missing. Here we show that both control and FMRP-KO hiPSCs can be converted to functionally mature cortical neurons and cerebral organoids, and that these in vitro systems display a FXS phenotype and can be considered promising disease models to study FXS and to develop and test disease-modifying drugs. Specifically, using an isogenic FMRP-KO hiPSC line we obtained a 2D mixed cortical culture displaying: i) altered neuronal and glial gene expression with increased expression of the astrocyte marker GFAP; ii) increased network activity; iii) increased excitation/inhibition ratio. Moreover we report for the first time the generation of a 3D brain organoid system to model FXS, characterized by: i) increased size ii) reduced neuronal staining and iii) increased GFAP immunoreactivity.

In our study, the time-series analysis revealed that the lack of FMRP increased the expression of some neural progenitor genes (*SOX2* and *FOXG1)* at day 10 and the levels of the astrocyte marker *GFAP*, with concomitant decrease of the neuronal precursor marker *TBR2*, at day 25. Hence, we speculate that FMRP is responsible for the correct balance and development of neuronal and glial components in cerebral cortex, in line with what observed in human neural progenitor cells [44] and in *Fmr1* KO mouse model [45].

One important point of strength of our system is that, differently to what observed in murine *Fmr1* KO cortical neurons [46], hiPSC derived FMRP-KO cultures are characterized by a gain of function of spontaneous network activity even in the absence of specific challenging protocols. Indeed, in control culture condition we observed that the silencing of FMRP induced a significant increase of both amplitude and frequency of “fast”, neuronal, spontaneous calcium events, and a significant increase of the amplitude of “slow” glial events.

Notably, together with the enhancement of network activity we report a different pattern of response to focal application of both glutamate and GABA in FMRP-KO cortical neurons, pointing to an unbalance of excitation and inhibition that relies on both systems. Indeed, while in our culture the large majority of cells responded to focal glutamate application with a fast intracellular calcium rise in both genotypes, the amplitude of glutamatergic evoked calcium transients was significantly higher in FMRP-KO neurons with respect to FMRP-WT. This phenomenon can be ascribed either to a functional upregulation of glutamatergic receptors, or to an increased depolarizing response, as reported for mouse neuronal progenitors lacking FMRP [47,48]. Moreover, although at day 70 the number of glutamatergic synapses was similar in FMRP-WT and FMRP-KO cortical cultures, as revealed by the pre (VGLUT1) and post-synaptic (PSD95) staining, we cannot exclude a different contribution of ionotropic and metabotropic glutamate receptors between the two genotypes in mediating the neuronal depolarization that might contribute to glutamate-induced calcium transients, as in murine neuronal progenitors [49].

Strikingly, we report here that in FMRP-KO culture the focal application of GABA induced a depolarizing response, as revealed by the appearance of a calcium transient evoked by GABA puff. This phenomenon may rely on an immature phenotype of FMRP-KO hiPSC-derived cortical neurons which could be characterized by a depolarized chloride equilibrium. Indeed, GABA, the major inhibitory neurotransmitter in the brain, binds to GABA-A (ionotropic) or GABA-B (metabotropic) receptors that in mature healthy neurons mediate an inhibitory response that leads to membrane hyperpolarization. GABA transmission plays indeed a key role in setting the excitation/inhibition ratio, which is abnormal in FXS, as well as in autism and other neurodevelopmental disorders [50]. Moreover, even if many studies into GABA-mediated mechanisms in FXS indicate a reduction in both GABA-A and GABA-B activation in *Fmr1*-KO mice [51–53], and in embryonic stem cell-derived human FXS neurons [54], GABA-A and GABA-B agonists that have been tried in clinical trials, failed in reducing cortical hyperexcitability [55].

We can speculate that this failure relies on the delayed developmental switch of chloride equilibrium in FXS; indeed, when the equilibrium potential for chloride is depolarized, GABA mediated responses depolarize neurons, increasing their excitability and triggering spiking and calcium entry into neurons as during development and in several pathological conditions [56–58]. Indeed, a delay in the switch of the chloride reversal potential has been reported in cortical neurons of *Fmr1* KO mice, causing the change in GABA transmission polarity, which in turn might affect neuronal development in the cortex of *Fmr1* KO mice [59]. In summary, we can speculate that, in FMRP-KO coltures, beside the unbalance of glutamatergic and GABAergic systems, the upregulation of astrocytes may contribute to the enhancement of network activity [60].

By taking advantage of hiPSC-derived cerebral organoids to model the complexity of human brain development, we were able to recapitulate some features of FXS. We found, in 3D cortical organoids derived from FMRP-KO hiPSCs, a significant increase in cortical organoid size, a reduction in neuronal marker MAP2 and an increase in astrocyte reactivity. The increase in GFAP staining was observed also in the 2D neural differentiation system and resembles what reported in primary glial cells isolated from *Fmr1* KO mice [45]. Astrocytes are well recognized as active players in the neurobiology of FXS and involved in the correct development and maturation of hippocampal synapses [61–63]. The presence of neuronal and glial cells in our FMRP-KO hiPSC derived 2D cultures and 3D organoids will thus provides an important tool for deepening our knowledge on astrocyte biology in FXS during the early stage of development.

We speculate that, in this novel 2D and 3D iPSC based models of human cortex, FMRP is responsible for the correct balance of neuron and glial development. This could be mediated, similarly to what observed in murine models, by the overexpression of GSK-3β [64] that, positively regulating Notch pathway [65], has been found to be sufficient to initiate a rapid and irreversible loss of neurogenesis by accelerating gliogenesis [66]. Thus we suggest that the 3D human brain organoid could efficiently recapitulate the in vivo pathological phenotypes.

In summary, we used the 3D organoid platform to recapitulate the development of cortical phenotype of FXS. This experimental platform can be potentially applied for FXS in vitro modelling and exploring effective treatments for this neurodevelopmental disorder.

## Author Contributions

Conceptualization, A.Ro., S.D.A., A.S. and A.Re.; Formal analysis, A.S., C.B., V.d.T., S.G. and M.R.; Investigation, C.B., F.S., A.S., F.C., V.d.T. ; Methodology, C.B., F.S., A.S., P.F.P., M.R. and F.C.; Project administration, A.Ro., S.D.A. and A.Re.; Supervision, S.D.A. and A.Ro.; Writing – original draft, S.D.A. and A.Ro.

## SUPPLEMENTARY MATERIAL

**Supplementary Figure S1.**
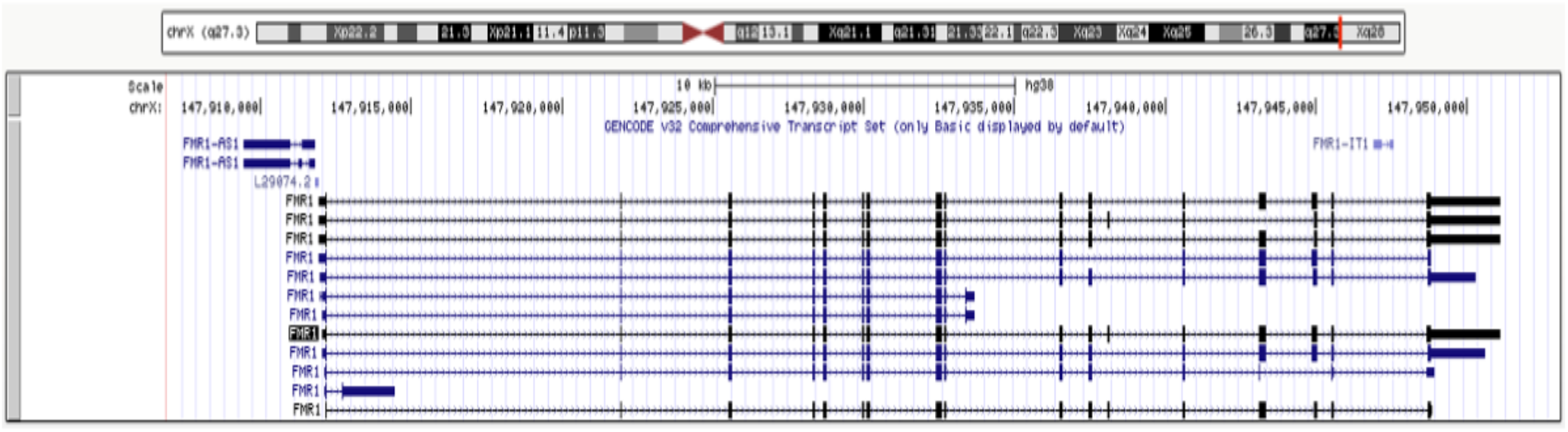
(A) Screenshot of the UCSC genome browser (http://genome.ucsc.edu; UCSC Genome Browser assembly ID: hg38) showing the human *FMR1* locus and the gene models included in the GENCODE track (v32 release). The track includes both protein-coding genes and non-coding RNA genes.

**Supplementary Figure S2.**
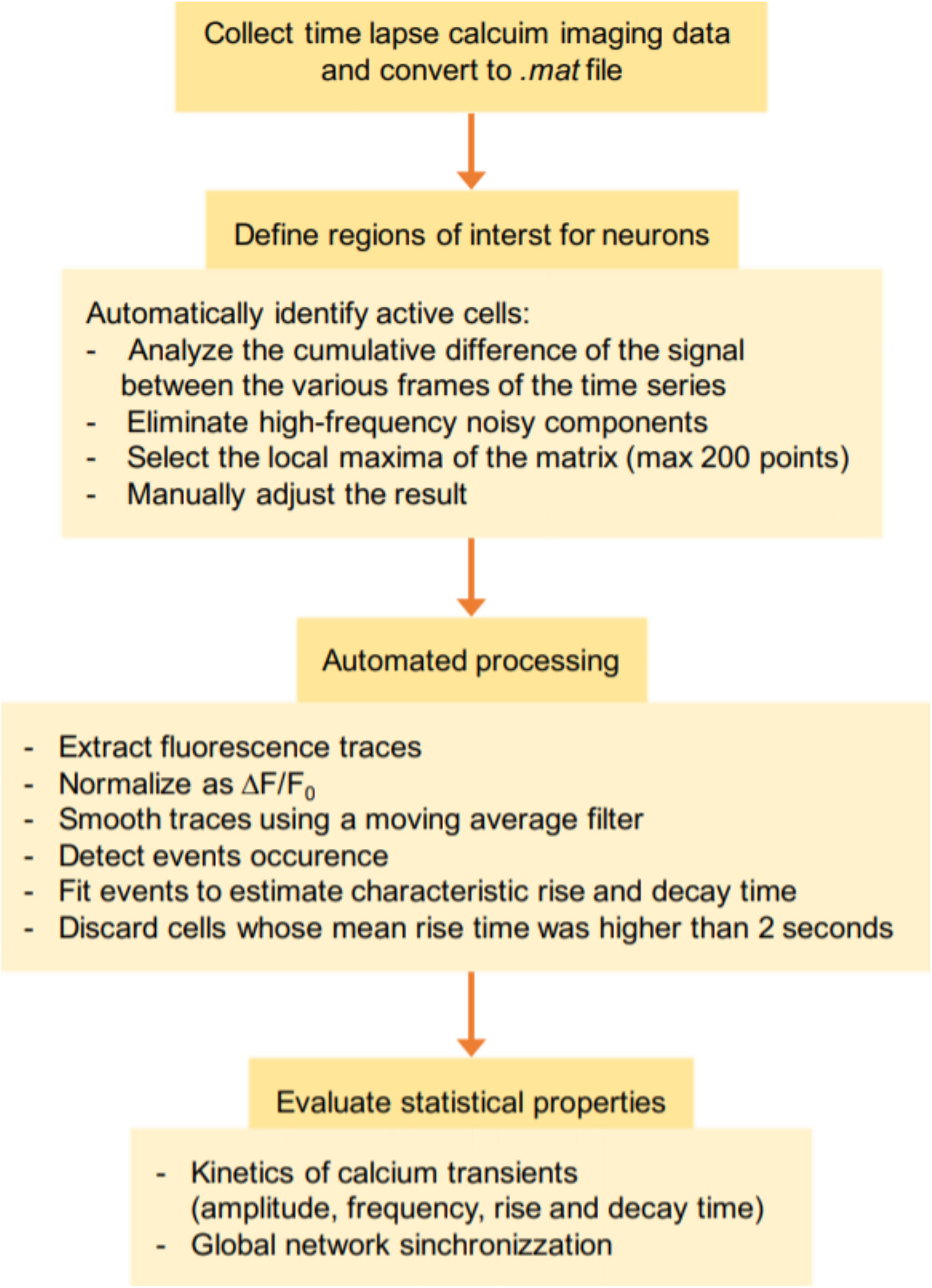
Schematic illustration of calcium imaging data processing performed in Matlab environment: data collected were analyzed through a custom-made algorithm designed to recognize cells, sort calcium events and select neuronal traces discarding traces characterized by slow rise time typical of astrocytes. Firing rate, amplitude and synchrony of the network were exported in Microsoft Excel to perform final statistics.

**Supplementary Figure S3.**
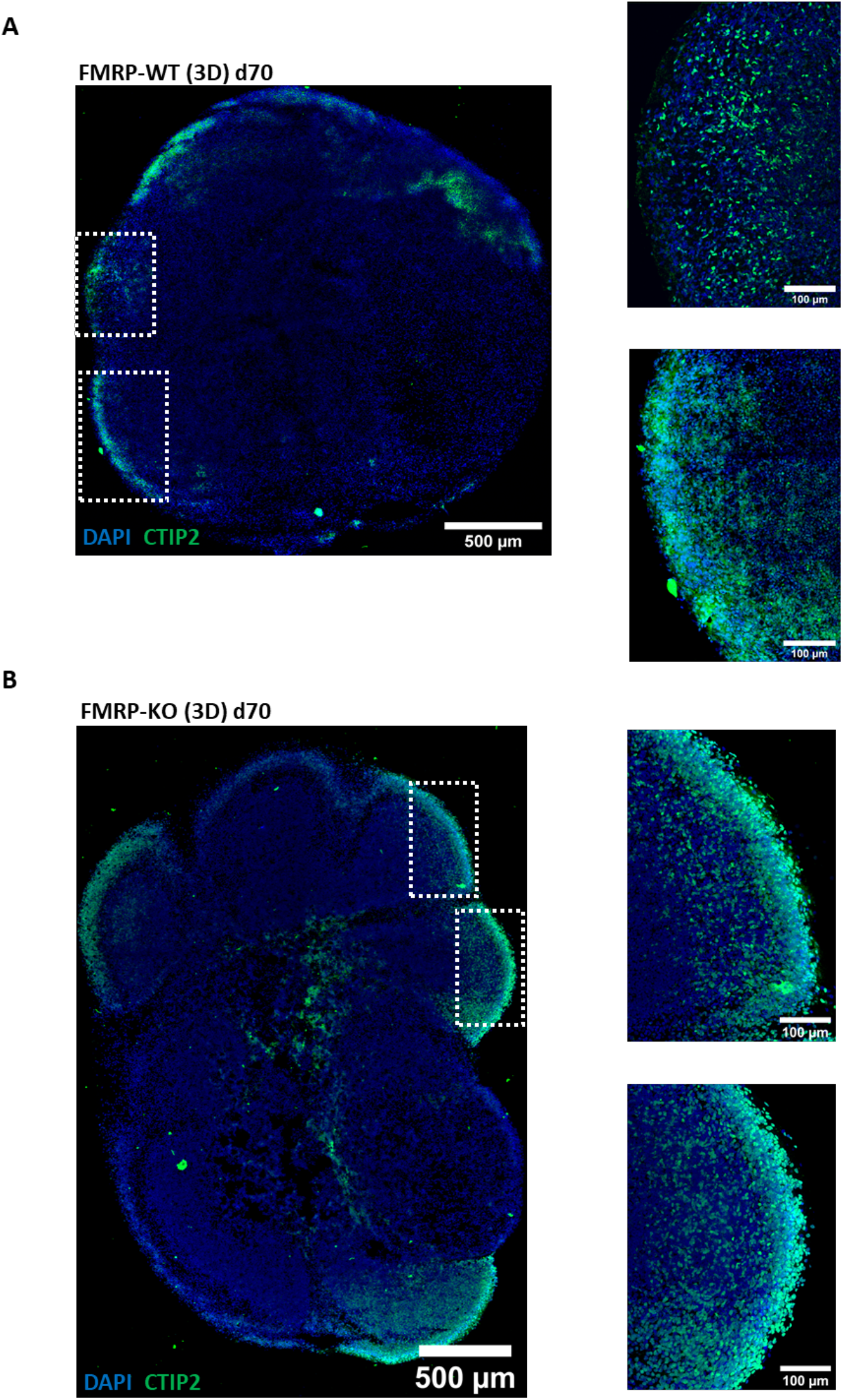
Cellular identity characterization revelead CTIP2 positive staining (green) indicating the presence of layer V cortical neurons at D70 of differentiation in both (A) FMRP-WT and (B) FMRP-KO brain organoids. Nuclei were stained with DAPI (blue). Scale bar: 500 μm. Insets show a close-up view of representative cortical plates. Scale bar: 100 μm.

**Supplementary Table 1.** List of secondary antibodies used for immunostaining in this study

| ***Antibody*** | ***Source*** | ***Product number*** | ***Supplier*** |
| --- | --- | --- | --- |
| Anti-mouse Alexa Fluor® 488 | goat | A11001 | Invitrogen |
| Anti-rat Alexa Fluor® 488 | goat | A11006 | Invitrogen |
| Anti-chicken Alexa Fluor® 488 | goat | A11039 | Invitrogen |
| Anti-rabbit Alexa Fluor® 488 | goat | A11008 | Invitrogen |
| Ancti chickenAlexa Fluor 594 | goat | A11042 | Invitrogen |
| Anti- mouseAlexa Fluor® 594 | goat | A11032 | Invitrogen |
| Anti-rabbit Alexa Fluor® 633 | goat | A21071 | Invitrogen |

**Supplementary Table 2.**
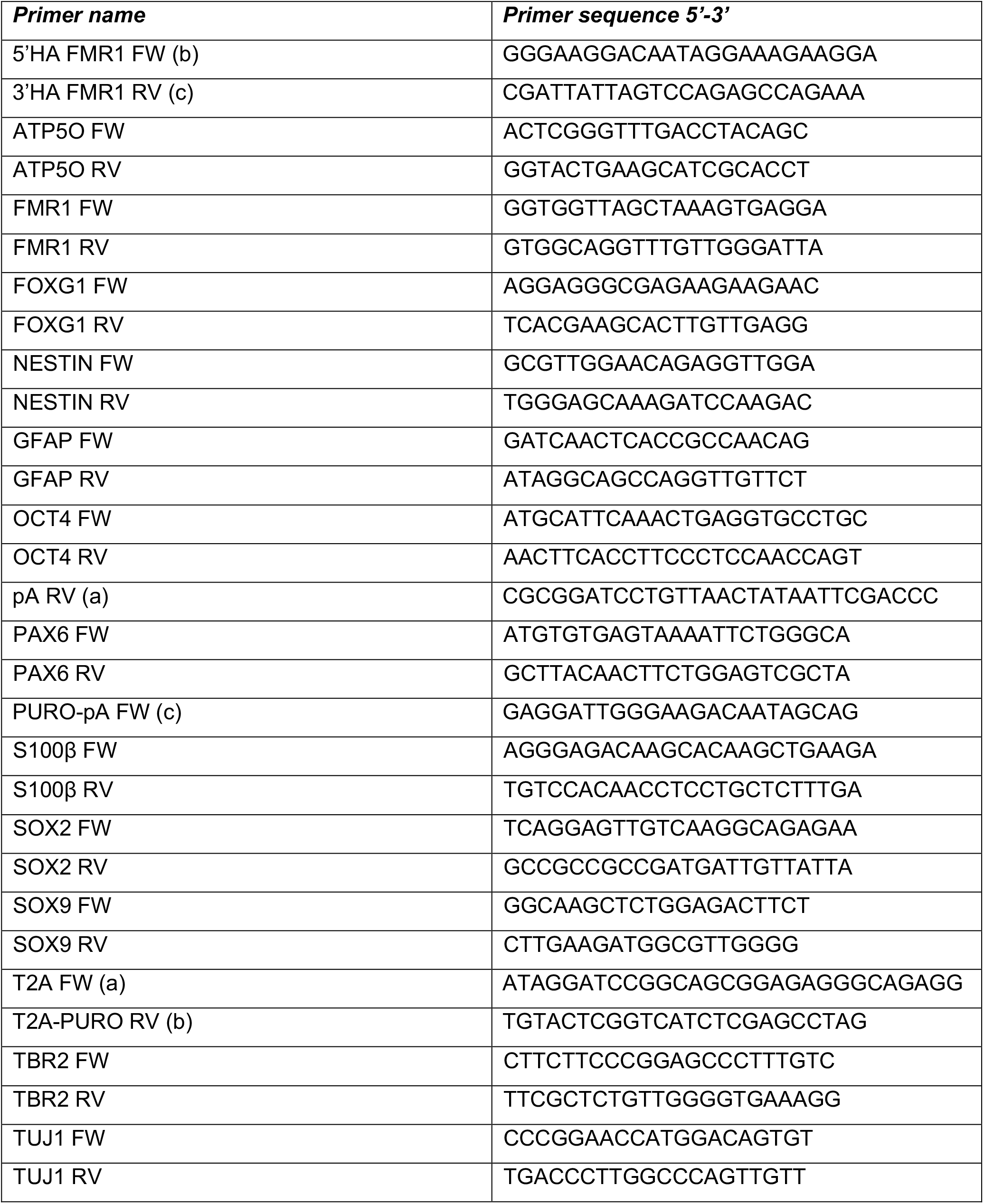
List of primers used in this study

